# Polycomb requires Tcp-1η chaperonin for maintaining gene silencing in *Drosophila*

**DOI:** 10.1101/2021.04.23.441114

**Authors:** Najma Shaheen, Jawad Akhtar, Zain Umer, Muhammad Haider Farooq Khan, Murtaza Saleem, Amir Faisal, Muhammad Tariq

**Affiliations:** Epigenetics and Gene Regulation Laboratory, Department of Biology, Syed Babar Ali School of Science and Engineering, Lahore University of Management Sciences, Lahore 54792, Pakistan; Department of Medicine, Division of Hematology/Oncology, Northwestern University Feinberg School of Medicine, Chicago, IL, 60611, USA; Department of Physics, Syed Babar Ali School of Science and Engineering, Lahore University of Management Sciences, Lahore 54792, Pakistan; Cancer Therapeutics Laboratory, Department of Biology, Syed Babar Ali School of Science and Engineering, Lahore University of Management Sciences, Lahore 54792, Pakistan

**Keywords:** Polycomb, Trithorax, TRiC, Tcp-1η, chaperonin, Cell fates, Epigenetic cellular memory

## Abstract

In metazoans, heritable states of cell type specific gene expression patterns linked with specialization of various cell types constitute transcriptional cellular memory. Evolutionarily conserved Polycomb group (PcG) and trithorax group (trxG) proteins contribute to the transcriptional cellular memory by maintaining heritable patterns of repressed and active expression states, respectively. Although chromatin structure and modifications appear to play a fundamental role in maintenance of repression by PcG, the precise targeting mechanism and the specificity factors that bind PcG complexes to a defined region in chromosomes remain elusive. Here we report a serendipitous discovery that uncovers a direct molecular interaction between Polycomb (PC) and TCP-1 Ring Complex (TRiC) chaperonin subunit, Tcp-1η in *Drosophila*. Tcp-1η interacts with PC at chromatin to maintain repressed states of homeotic and non-homeotic targets of PcG, which supports a strong genetic interaction observed between *Pc* and *Tcp-1η* mutants. Depletion of Tcp-1η results in dissociation of PC from chromatin and redistribution of an abundant amount of PC in cytoplasm. We propose that Tcp-1η is an important modulator of PC, which helps PC recruitment at chromatin and compromising Tcp-1η can directly influence an evolutionary highly conserved epigenetic network that supervises the appropriate cellular identities during development and homoeostasis of an organism.

**Significance Statement:** Silencing of key developmental genes, e.g, Hox genes, by PcG is a hallmark of differential gene expression patterns associated with cell fate determination. Here we describe a previously unknown molecular and genetic interaction of Polycomb (PC) with Tcp-1η subunit of TRiC chaperonin complex in *Drosophila*. Compromising Tcp-1η function results in de-repression of PcG targets and a concomitant loss of PC from chromatin. Moreover, depletion of Tcp-1η leads to redistribution of PC in cytoplasm. Molecular interaction of PC with Tcp-1η highlights a novel factor which helps PC recruitment at chromatin. We propose that Tcp-1η chaperonin is one of the specificity factors and part of the targeting mechanism that binds PC to specific regions on chromosomes.

## Introduction

In multicellular eukaryotes, epigenetic factors play a pivotal role in cell fate determination visualized by the differential gene expression profiles established during the patterning process in early development. In particular, maintenance of transcriptional states of important regulators like the Hox genes ensures faithful propagation of determined states during cell proliferation (1, 2). The Polycomb group (PcG) and trithorax group (trxG) proteins maintain the heritable patterns of repressed and active gene expression states, respectively. An evolutionarily conserved hallmark of PcG and trxG regulation is the establishment and maintenance of long-term developmental decisions. Both groups exert their functions by interacting with the histones, the transcription machinery and by generating heritable epigenetic marks on chromatin (3, 4).

Molecular and biochemical analyses in *Drosophila* have unraveled the composition of at least two different PcG complexes, i.e., PRC1 and PRC2, and many accessory factors. Silencing by the PcG complexes correlates with histone H3 lysine 27 tri-methylation (H3K27me3) (5–7) and histone H2A lysine 118 ubiquitination (H2AK118ub) (8). On the other hand, trxG proteins, discovered as suppressors of PcG, act as anti-silencing factors (9) and several of them are known to act by modifying local properties of chromatin. For example, TRX and ASH1 directly counteract repression by catalyzing tri-methylation of histone H3 lysine 4 (H3K4me3) and histone H3 lysine 36 (H3K36me3), respectively. Moreover, trxG is much more heterogeneous with respect to protein complexes that covalently modify chromatin, remodel the chromatin and interact with general transcription machinery (3). Molecular chaperones like Hsc4 (Heat shock cognate 4) (10), Hsp90 (Heat shock protein 90) (11, 12) and immunophilins (13–15) are proposed to serve as additional factors for PcG/trxG to maintain heritable patterns of gene expression. For example, the *Hsc4* mutant flies exhibit *Pc* like phenotype and Hsc4 was found to be part of the PcG multi-protein complex (10). Besides Hsc4, a novel *Drosophila* J class chaperone (Droj2) was also found stably associated with Polyhomeotic, a PcG protein, in *Drosophila* (16). Additionally, mutations in Hsp90 mimic trxG like behavior and TRX requires Hsp90 for maintenance of gene activation in *Drosophila* (11). Genome-wide binding profile of Hsp90 shows a massive overlap with TRX at chromatin. Notably, Hsp90 also contributes to paused RNA polymerase and facilitates transcriptional block by PcG (12). The presence of RNA polymerase and general transcription factors at silent promoters (9, 17–19) and the competition observed between PcG and trxG to regulate the gene expression suggest that PcG and trxG proteins associate with their target genes as dynamic complexes which are in balance with one another (20). However, the mechanisms and the factors responsible to shift the balance in favor of either PcG or trxG during their dynamic association with chromatin remain elusive.

Here we report a serendipitous discovery that describes direct molecular interactions of *Drosophila* PC with Tcp-1η subunit of TCP-1 ring complex (TRiC), a class of chaperones also called chaperonins. The TRiC chaperonins are protein-folding machines comprising of hetero-oligomeric, double ringed, high molecular weight, ATP dependent chaperones involved in the folding and assembly of multi-protein complexes (21). The complex architecture and mechanism of action of TRiC allow it to chaperone essential proteins involved in diverse cellular processes like cell cycle regulation (22–24), cytoskeletal organization (25), organ size (26) and signal transduction pathways (27–30). In our quest for novel regulators of trxG, we had discovered *Drosophila Tcp-1η* and *CCT5* subunits of TRiC as top trxG candidates influencing luciferase reporter in an ex vivo genome-wide RNAi screen (31). However, contrary to our earlier discovery of *Tcp-1η* and *CCT5* as trxG candidates in the RNAi screen, further analysis revealed that both *Tcp-1η* and *CCT5* mutants genetically interact with *Polycomb* (*Pc*) and strongly enhance extra sex comb phenotype. Moreover, suppression of *trx* mutant phenotype by *Tcp-1η* mutation and a strong reactivation of homeotic genes in homozygous *Tcp-1η* mutants corroborate the PcG like behavior of these TRiC subunits. Importantly, presence of Tcp-1η together with PC on polytene chromosomes and its association at PcG targets, as characterized by chromatin immunoprecipitation, support our hypothesis that Tcp-1η indeed contributes to repression by PcG. Our results further demonstrate that depletion of Tcp-1η results in dissociation of PC from the chromatin, which explains the reactivation of PcG targets upon *Tcp-1η* knock-down. Notably, Tcp-1η was found to interact with PC at the molecular level in *Drosophila* cells and a substantial amount of PC was found in the cytoplasm when Tcp-1η was depleted. Together our data provides an affirmative evidence for the role of Tcp-1η in regulating PcG mediated gene expression patterns.

## Results

### TCP-1 Ring Complex (TRiC) genetically interacts with PcG and trxG system

Since *Tcp-1η* and *CCT5* subunits of TRiC chaperonin complex emerged as candidate trxG genes in a luciferase reporter based genome-wide RNAi screen in *Drosophila*, we started validation of *Tcp-1η* and *CCT5* as potential trxG genes. To investigate if TRiC subunits genetically interact with PcG/trxG system, mutants for *Tcp-1η* (*Tcp-1η*^*KG01477*^ and *Tcp-1η*^*KG09501*^) and *CCT5* (*CCT5*^*06444*^ and *CCT5*^*K06005*^) were crossed with two different *Pc* (*Pc*^*1*^ and *Pc*^*XL5*^) alleles and males from F1 progeny (*Tcp-1η****/****Pc* and *CCT5/+;Pc/+*) were analyzed for extra sex comb phenotype. Heterozygous *Pc* (*+/Pc*^*1*^ and *+/Pc*^*XL5*^) male flies, obtained from a cross of *Pc* (*Pc*^*1*^ and *Pc*^*XL5*^) mutants with *w*^*1118*^ mutant flies, exhibit strong extra sex comb phenotype. As compared to heterozygous *Pc* mutant males, used as control, each mutant allele of *Tcp-1η* and *CCT5* significantly enhanced extra sex comb phenotype of *Pc*^*1*^ and *Pc*^*XL5*^ (Fig. 1 *A-D* and *SI Appendix*, Fig. S1 *A-D*). This strong genetic interaction of two different alleles of *Tcp-1η* as well as *CCT5* with both the alleles of *Pc* indicates that *Tcp-1η* and *CCT5* mutants behave like *PcG* genes. However, these results were contrary to the discovery of *Tcp-1η* and *CCT5* as candidate trxG genes in the genome-wide RNAi screen. To further validate PcG like behavior of TRiC subunits, it was investigated if *Tcp-1η* mutants antagonize *trx* mutant phenotype which is a hallmark of PcG genes. Both the mutant alleles of *Tcp-1η* were crossed with *trx* (*trx*^*1*^) mutant to investigate if *Tcp-1η* genetically interacts with *trx* and suppresses A5 to A4 homeotic phenotype of *trx* mutants. Heterozygous *trx* (*trx/+*) mutant males from the cross of *w*^*1118*^ with *trx* exhibit loss of abdominal pigmentation (Fig. 1 *E*, right), classified as A5 to A4 transformation (32). Both the alleles of *Tcp-1η* strongly suppressed A5 to A4 transformation (Fig. 1 *E*, left) in *Tcp-1η/trx* double mutants and resulted in a higher percentage of male flies with wild type abdominal pigment (Fig. 1 *F*). A strong suppression of *trx* phenotype as well as enhancement of extra sex comb phenotype of *Pc* by both alleles of *Tcp-1η* support the notion that *Tcp-1η* acts as a PcG gene. Since luciferase is a known client of TRiC chaperonin (33), it explains the presence of *Tcp-1η* and *CCT5* subunits as trxG candidates in luciferase based genome-wide RNAi screen.

**Fig. 1.**
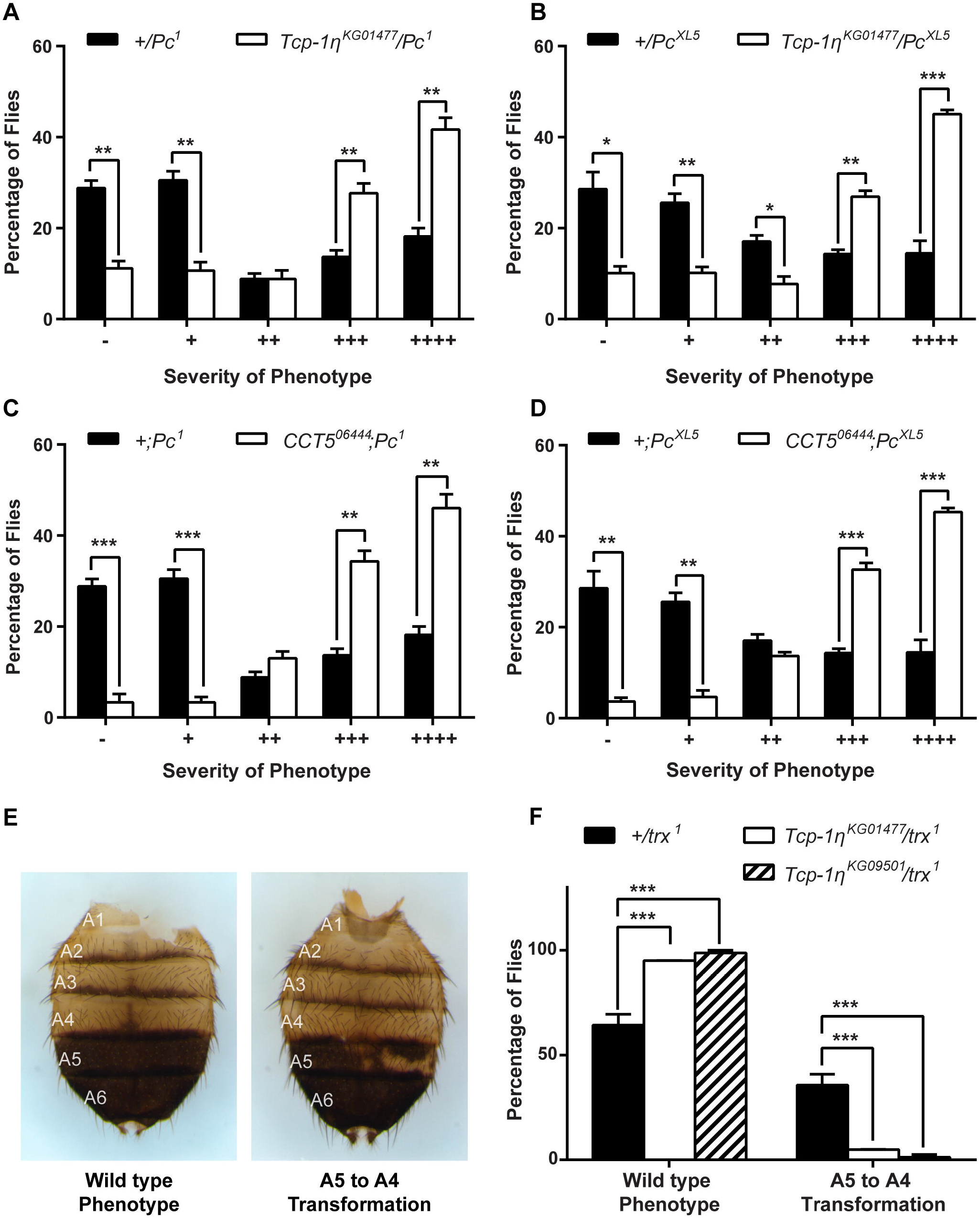
Mutants of TRiC exhibit PcG like behavior. (A-D) Heterozygous male flies for *Pc* (*+/Pc*^*1*^ and *+/Pc*^*XL5*^) obtained by crossing *Pc* (*Pc*^*1*^ and *Pc*^*XL5*^) alleles with *w*^*1118*^ exhibit strong extra sex comb phenotype and these heterozygous males were used as control. *Tcp-1η* (*Tcp-1η*^*KG01477*^) and *CCT5* (*CCT5*^*06444*^) mutants crossed to two different alleles of *Pc* (*Pc*^*1*^ and *Pc*^*XL5*^) significantly enhanced extra sex comb phenotype in *Tcp-1η*^*KG01477*^*/Pc* (A, B) and *CCT5*^*06444*^*;Pc* (C, D) mutants, respectively, when compared with control (*+/Pc*^*1*^ and *+/Pc*^*XL5*^). More than 150 male flies with desired genotype from the progeny of each cross were scored for extra sex combs. Flies were categorized based on the number of extra sex comb bristles on the second and third pair of legs (11) as follows: −; no extra sex combs on second or third pair of legs, +; 1–2 bristles on the second leg, ++; 3 or more bristles on the second leg, +++; 3 or more bristles on the second leg and 1–2 bristles on the third leg and ++++; strong sex combs on both second and third pairs of legs. The percentage of flies for each category was plotted as bar graphs. Error bars represent standard error means (SEM) from three independent experiments. Statistical significance was calculated using t-test (* p ≤ 0.05, ** p ≤ 0.01, *** p ≤ 0.001, **** p ≤ 0.0001). (E, F) *Tcp-1η/trx*^*1*^ double mutant males from progeny of *Tcp-1η* mutants crossed to *trx*^*1*^ were scored for suppression of A5 to A4 transformation. Heterozygous *trx* mutant (*+/trx*^*1*^) males obtained from the cross of *w*^*1118*^ with *trx*^*1*^ mutant exhibit abdominal pigment phenotype referred to as A5 to A4 transformation (E, right) and were used as controls. (F) *Tcp-1η* mutations suppressed *trx* mutant phenotypes in double mutant *Tcp-1η/trx*^*1*^ male flies as compared to control heterozygous male flies for *trx* (*+/trx*^*1*^). Flies were categorized into wild type (E, left) and A5 to A4 transformation (E, right) based on abdominal pigmentation phenotype. Percentage of flies obtained for each category of phenotype was plotted as bar graphs (F). Error bars represent SEM from two independent experiments. Statistical significance was calculated using two-way ANOVA (* p ≤ 0.05, *** p ≤ 0.001, **** p ≤ 0.0001).

### *Tcp-1η* is required for PcG mediated gene silencing

Since a strong genetic interaction of *Tcp-1η* with PcG system was observed, we aimed to investigate if *Tcp-1η* contributes to maintenance of repression by PcG. To this end, 12hrs old homozygous *Tcp-1η* mutant embryos were stained with Abd-B antibody and compared with *w*^*1118*^ embryos of the same age, which were used as control (Fig. 2 *A-I*). In *w*^*1118*^ embryos at this stage, *Abd-B* is expressed at progressively increasing level from parasegment 10 (PS10) to PS14 exhibiting strongest expression in PS14 (34, 35) (Fig. 2*C*). As compared to *w*^*1118*^ control, *Tcp-1η* mutant embryos showed a drastic increase in Abd-B expression (Fig. 2 *F* and *I*). This increased Abd-B expression is a hallmark of PcG mutants (36). Additionally, *Tcp-1η* was knocked down in D.Mel-2 cells using dsRNA (Fig. 2*J*) and expression of PcG target genes was analyzed using quantitative real-time PCR. As compared to *LacZ* dsRNA treated cells used as control, depletion of Tcp-1η resulted in a significant upregulation of *psq* (*pipsqueak*), *Antp* (*Antennapedia*), *Abd-B* (*Abdominal B*), *Dfd* (*Deformed*) and *Ubx* (*Ultrabithorax*) (Fig. 2*K*) which are known PcG targets (19, 37, 38). Next, *Tcp-1η* was knocked down in flies using eye-specific GAL4 driver line (Fig. 2 *L-N*). Depletion of Tcp-1η resulted in transformation of eye to duplicated antenna (Fig. 2*M*) and appearance of a leg-like appendage besides reduced eye size (Fig. 2*N*). These phenotypes and the headless pupal lethal pharate adults observed upon depletion of Tcp-1η (*SI Appendix*, Fig. S2*C*) may be attributed to the ectopic expression of *Antp* in eye-antennal imaginal discs (39–41). Similar pupal lethal phenotypes have been reported in eye-specific knock-down of all eight subunits of TRiC chaperonin complex (26). Together, these results suggest that PcG requires Tcp-1η to maintain repression.

**Fig. 2.**
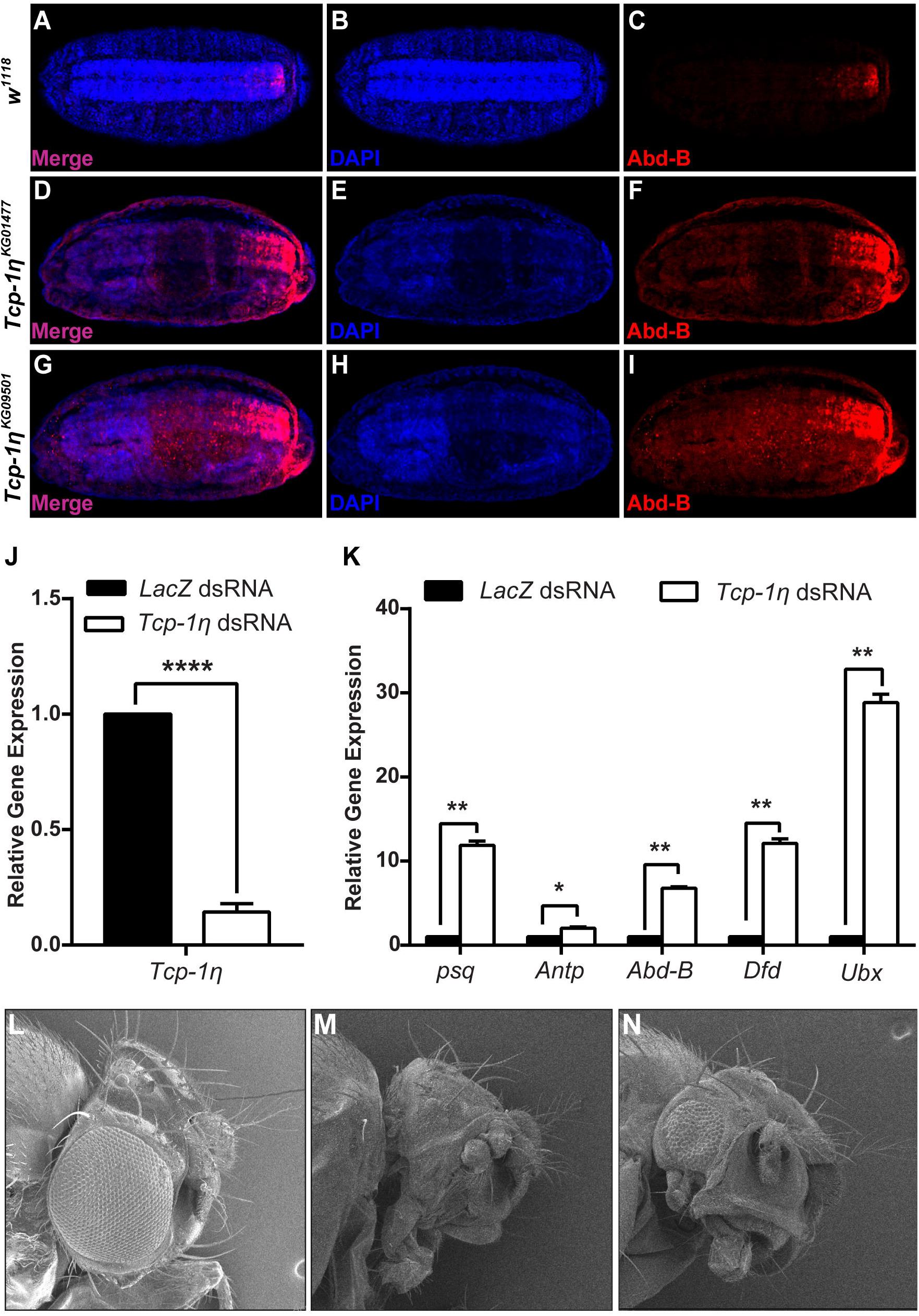
Depletion of Tcp-1η enhances the expression of PcG target genes. (A-I) Enhanced levels of Abd-B expression observed in *Tcp-1η* mutants compared with *w*^*1118*^. Stage 15 embryos of *w*^*1118*^ (A-C), homozygous mutant embryos for *Tcp-1η*^*KG01477*^ (D-F) and *Tcp-1η*^*KG09501*^ (G-I) were stained with Abd-B antibody. Homozygous *Tcp-1η* embryos (F, I) showed increased Abd-B expression as compared to *w*^*1118*^ embryos (C) used as control. All embryos are oriented with their anterior ends to the left. (J) As compared to *LacZ* dsRNA treated cells, knock-down of *Tcp-1η* showed drastic reduction in mRNA levels of *Tcp-1η* in D.Mel-2 cells. (K) Significantly increased levels of *psq, Antp, Abd-B, Dfd* and *Ubx* expression determined in *Tcp-1η* knock-down cells (J) using real-time PCR compared with *LacZ* dsRNA treated cells. Error bars represent SEM for two independent experiments. Statistical significance was calculated using t test (* p ≤ 0.05, ** p ≤ 0.01, *** p ≤ 0.001, **** p ≤ 0.0001). (L-N) Eye specific knock-down of *Tcp-1η* shows a range of homeotic phenotypes, i.e., transformation of eye to duplicated antenna (M) and appearance of a leg-like appendage in addition to reduction in eye size (N).

### Tcp-1η binds to chromatin at PcG targets

Since homozygous mutation of *Tcp-1η* in vivo and its depletion in cells result in de-repression of PcG targets, it was questioned if Tcp-1η associates with chromatin at PcG targets and contributes to maintenance of repression. To address this question, transgenic flies expressing *Myc-Tcp-1η* (*SI Appendix*, Fig. S2*A*) under GAL4 inducible promoter were generated. Polytene chromosomes were prepared from third instar larvae expressing *Myc-Tcp-1η* in salivary glands and stained with anti-Myc and anti-PC antibodies (Fig. 3 *A-D*). It was observed that Tcp-1η co-localizes with PC at multiple sites on polytene chromosomes (Fig. 3*D*). Next, chromatin immunoprecipitation (ChIP) was performed from *Drosophila* S2 stable cells expressing *FLAG-Tcp-1η* using anti-FLAG antibody (*SI Appendix*, Fig. S2*B*). ChIP from empty vector (*EV*) control cells expressing FLAG-epitope under copper inducible promoter was used as a control. As compared to ChIP from *EV* control cells, FLAG-Tcp-1η was found to be enriched at *psq, Dfd* and *bxd* (*bithoraxoid*) (Fig. 3*E*) which are known targets of PcG (19, 37, 38). Importantly, Tcp-1η was not present at an intergenic region (IR) used as control where PC does not bind (Fig. 3*E*). Based on the chromatin association of Tcp-1η at PcG target genes, it was assumed that Tcp-1η may biochemically interact with PC and facilitate maintenance of silencing by PcG. To investigate molecular interaction between Tcp-1η and PC, co-immunoprecipitation (Co-IP) was performed from stable cells containing *FLAG-Tcp-1η* transgene. Total cell lysates from *EV* control and *FLAG-Tcp-1η* stable cells were subjected to pull down with anti-PC antibody and subsequently analyzed on a Western blot which was probed with anti-FLAG and anti-PC antibodies. As compared to Co-IP from *EV* control, immunoprecipitation with anti-PC specifically resulted in enrichment of FLAG-Tcp-1η from stable cells expressing FLAG-Tcp-1η (Fig. 3*F*), indicating that Tcp-1η interacts with PC in *Drosophila* cells. The chromatin association of Tcp-1η on PcG target genes and its interaction with PC signify a close interaction between the two to maintain gene silencing.

**Fig. 3.**
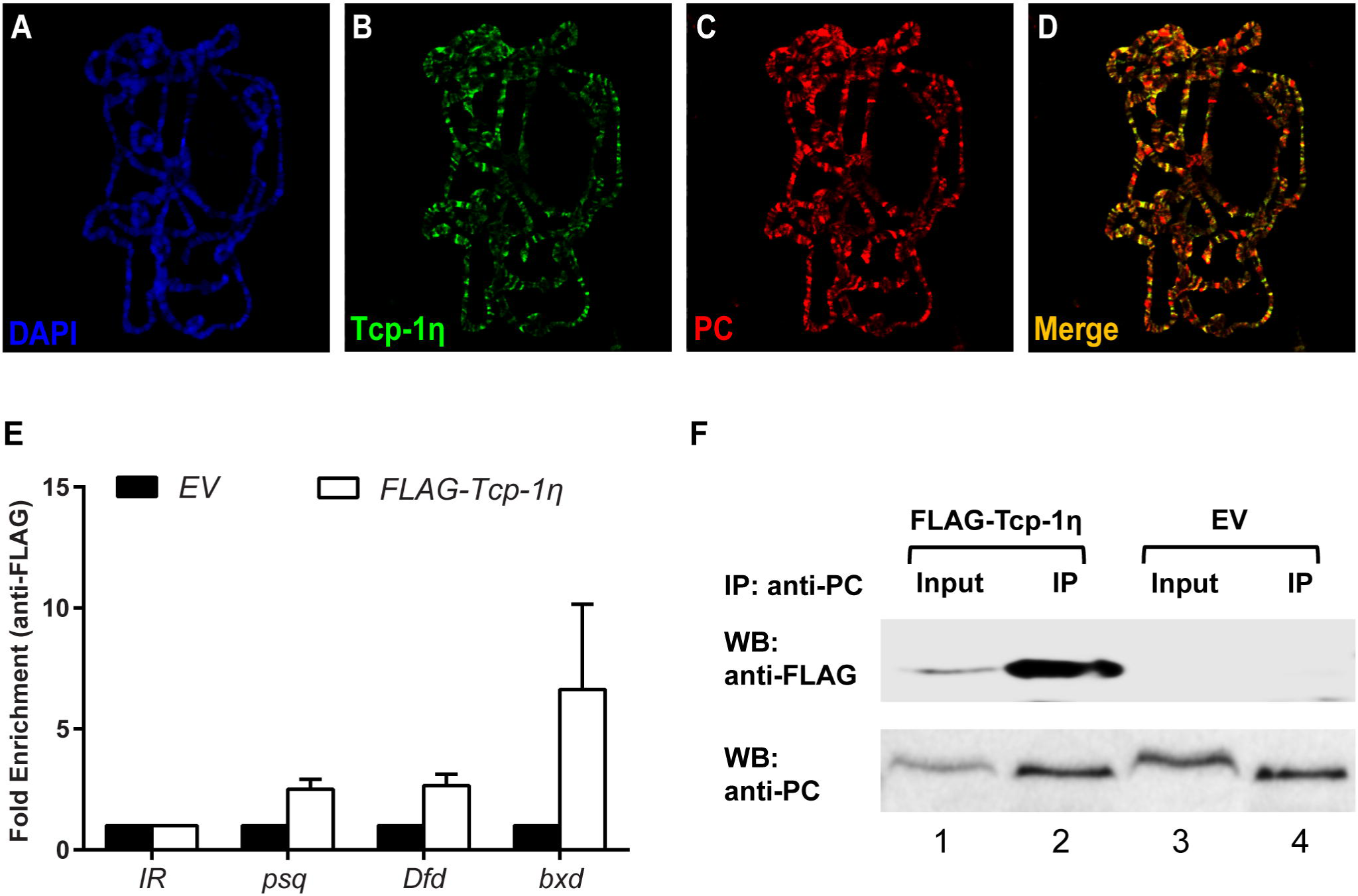
Tcp-1η co-localizes and interacts with PC. (A-D) Polytene chromosomes prepared from third instar larvae expressing Myc-Tcp-1η were stained with anti-Myc (B) and anti-PC (C) antibodies. Myc-Tcp-1η was observed to co-localize with PC at numerous loci, seen as yellow bands in merge (D). (E) ChIP performed from FLAG-Tcp-1η expressing cells using anti-FLAG antibody showed Tcp-1η enriched at known PcG target genes, i.e., *psq, Dfd* and *bxd* as compared to *EV* (empty vector) control cells. *IR* (Intergenic region) represents region where PC is normally not bound (59). ChIP was analyzed using fold enrichment over *IR*. Error bars represent SEM for two independent experiments. (F) PC biochemically interacts with Tcp-1η. Co-Immunoprecipitation (Co-IP) was performed using anti-PC antibody from *FLAG-Tcp-1η* expressing stable cells as well as empty vector (*EV*) control cells. Western blot with anti-FLAG (WB: anti-FLAG) showed specific enrichment of FLAG-Tcp-1η in the IP sample (lane 2) of *FLAG-Tcp-1η* expressing cells as compared to control IP from *EV* cells (lane 4). Western blot with anti-PC (WB: anti-PC) showed specific and equal enrichment of PC in IP from both *FLAG-Tcp-1η* expressing cells as well as *EV* cells. Input represents 1% of total cell lysates from *FLAG-Tcp-1η* (lane 1) and *EV* (lane 3) cells.

### Tcp-1η facilitates nuclear localization and chromatin association of PC

The genetic and molecular interaction of Tcp-1η with PC intrigued us to ask if depletion of Tcp-1η has an impact on association of PC at the chromatin. To address this question, ChIP was performed using anti-PC antibody from cells treated with dsRNA against *Tcp-1η*. As compared to control cells treated with *LacZ* dsRNA, ChIP from Tcp-1η depleted cells revealed reduced association of PC at *bxd* and *Dfd* chromatin binding sites (Fig. 4*A*). Moreover, ChIP analysis of RNA pol-II binding in *Tcp-1η* knock-down cells revealed that RNA pol-II was enriched along the gene body of *Dfd* (Fig. 4*B*) which correlates with increased expression of *Dfd* in these cells (Fig. 2*K*). For an in vivo validation of decreased association of PC at chromatin after Tcp-1η depletion in cells, *Tcp-1η* was knocked down specifically in salivary glands of third instar larvae by crossing *Tcp-1η* RNAi flies with salivary gland specific GAL4 (*Sgs-GAL4*) driver line (Fig. 4 *C-H*). Polytene chromosomes were prepared from third instar larvae and immunostaining was performed with PC as well as RNA pol-II antibodies. Polytene chromosomes from *w*^*1118*^ larvae of the same age stained with PC and RNA pol-II antibodies were used as control (Fig. 4 *C-E*). As compared to control (Fig. 4*E*), PC staining was drastically reduced on polytene chromosomes from salivary glands where Tcp-1η was depleted (Fig. 4*H*). However, no such effect on RNA pol-II association with polytene chromosomes was observed in Tcp-1η depleted salivary glands when compared with polytenes from *w*^*1118*^ control (Fig. 4 *D* and *G*). This PC specific effect of Tcp-1η depletion led us to investigate possible underlying effects on the expression of *Pc*. To this end, we knocked down *Tcp-1η* using dsRNA and analyzed the expression of *Pc*.

**Fig. 4.**
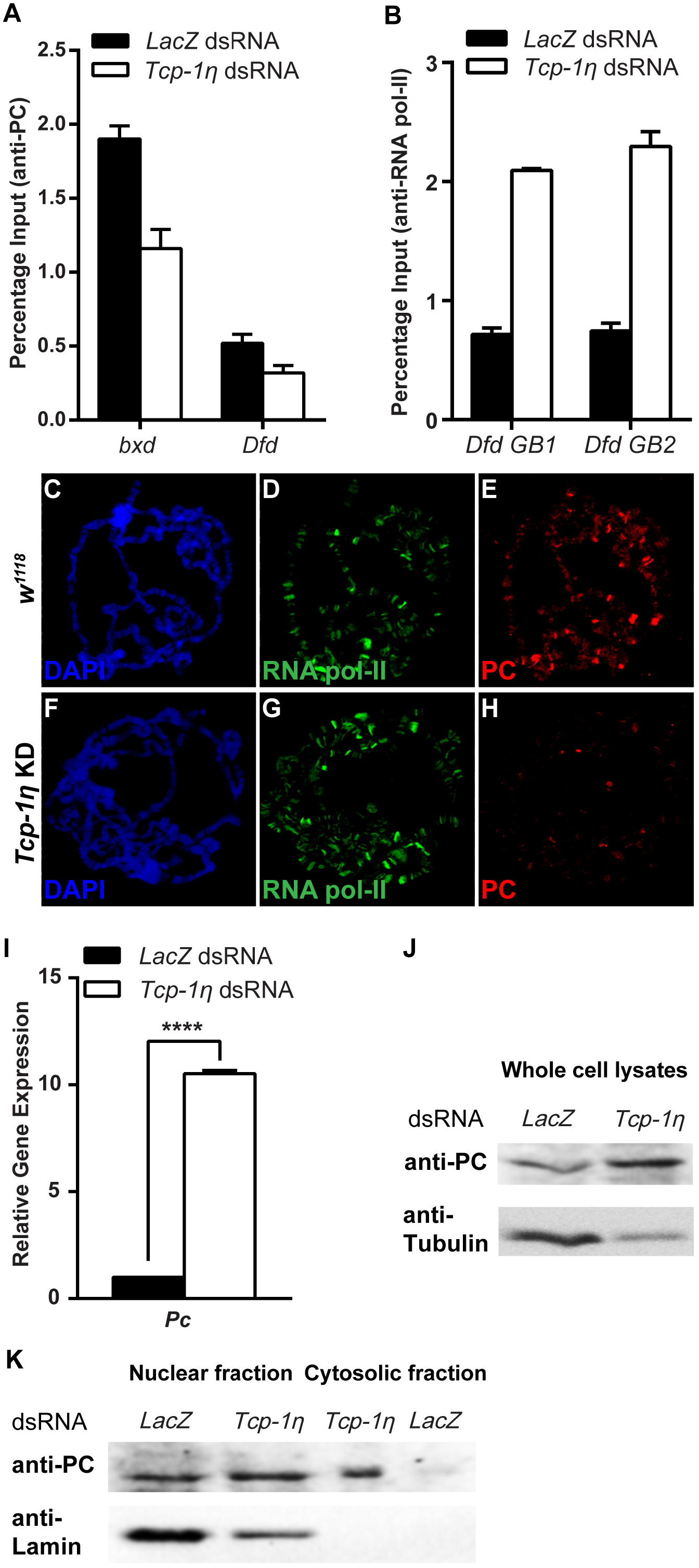
Tcp-1η is required for association of PC at chromatin. (A) ChIP using anti-PC antibody from *Tcp-1η* knock-down cells showed decreased enrichment of PC at *bxd* and *Dfd* as compared to *LacZ* dsRNA treated control cells. (B) ChIP from cells treated with dsRNA against *Tcp-1η* showed increased RNA pol-II along the gene body of *Dfd* (labelled *Dfd GB1* and *Dfd GB2*) as compared to control correlating with an increased expression of *Dfd* (Fig. 2*K*) in Tcp-1η depleted cells. (C-H) Polytene chromosomes prepared from *w*^*1118*^ and Tcp-1η depleted salivary glands and stained with anti-RNA pol-II and anti-PC antibodies. As compared to RNA pol-II staining used as a positive control (D, G), a strongly diminished binding of PC was observed after depletion of Tcp-1η (H) when compared with *w*^*1118*^ control (E). (I, J) Drastic increase in expression of *Pc* in Tcp-1η depleted cells. Tcp-1η depleted cells displayed significantly higher expression of *Pc* in real time PCR analysis (I) as well as on Western blot (J) when compared to control cells. Error bars represent SEM for two independent experiments. Statistical significance was calculated using t-test (**** p ≤ 0.0001). (K) Western blot probed with anti-PC and anti-Lamin antibodies showing nuclear and cytosolic fractions isolated from Tcp-1η depleted and control cells. Besides its presence in nucleus, PC was also found to be enriched in the cytosolic fraction of Tcp-1η depleted cells as compared to cytosolic fraction of *LacZ* dsRNA treated control. The same blot probed with anti-Lamin served as a control for purity of nuclear and cytosolic fractions.

Interestingly, quantitative real-time PCR analysis revealed that expression of *Pc* was significantly increased in Tcp-1η depleted cells as compared to *LacZ* dsRNA treated cells used as control (Fig. 4*I*). This increase in the mRNA levels of *Pc* also correlates with increased PC protein in *Tcp-1η* knock-down cells as compared to control cells (Fig. 4*J*). However, the increased expression of PC in cells is contrary to the decreased association of PC on chromatin when Tcp-1η was depleted (Fig. 4 *A* and *H*), which led us to investigate if Tcp-1η is required for proper localization of PC within the cells. To address this question, we isolated nuclear and cytosolic fractions of Tcp-1η depleted cells and analyzed the fractions on a Western blot with anti-PC antibody. Intriguingly, an increased amount of PC was observed in the cytosolic fraction of *Tcp-1η* knock-down cells when compared with control cells. However, amount of PC in the nuclear fractions of Tcp-1η depleted as well as control cells remained the same. Absence of Lamin in the cytosolic fraction served as control and illustrates that there was no nuclear contamination in the cytosolic fraction. Therefore, it validates that the PC observed in the cytosolic fraction is a specific consequence of Tcp-1η depletion (Fig. 4*K*). The increased expression of PC in Tcp-1η depleted cells does not get associated with chromatin suggesting that Tcp-1η is required for recruitment of PC at chromatin. Moreover, presence of an ample amount of PC in cytoplasm in the absence of Tcp-1η further suggests that Tcp-1η is more likely playing a role in chaperoning PC to its chromatin binding sites. Taken together, these results highlight a critical role for Tcp-1η in maintenance of gene repression by PC in the process of cell fate maintenance and epigenetic cell memory in *Drosophila*.

## Discussion

In order to understand better the complex relationship between genotype and phenotype it is necessary to study the link and interplay between epigenetic pathways, cellular signaling and environmental factors during development. Molecular chaperones represent a class of proteins, which rapidly respond to intracellular or extracellular environmental cues to initiate epigenetically heritable changes (42, 43). Our results demonstrate a previously unknown genetic and molecular link between Tcp-1η and PC which is essential for maintaining heritable gene repression. We provide evidence that depletion of Tcp-1η results in reactivation of PcG target genes and leads to severe morphological defects. Since PC dissociates from chromatin and is retained in cytoplasm as a consequence of diminished Tcp-1η, we propose that Tcp-1η is required for chaperoning PC to its targets on the chromatin and maintenance of repression. Involvement of Tcp-1η in maintenance of silencing by PC has revealed a novel epigenetic modulator of PC which ensures robustness of gene expression during development. Such an interaction of molecular chaperone Hsp90 with epigenetic factors like TRX has previously been reported in *Drosophila* (11, 12, 44). In contrast to the role of Hsp90 in gene activation by trxG, little is known about role of molecular chaperones in PcG mediated gene regulation and the effects that they may have on epigenetic cell memory.

Genetic evidence presented here demonstrates that TRiC complex members, *Tcp-1η* and *CCT5*, exhibit PcG like behavior by enhancing extra sex comb phenotype of *Pc*. Appearance of extra sex combs on the second and third pairs of legs in *Drosophila* males is a classical mutant phenotype in *Pc* heterozygotes (45, 46). Strong enhancement of extra sex comb phenotype by two different alleles of *Tcp-1η* as well as *CCT5* suggests that TRiC subunits are possibly required for the proper function of one or more PcG proteins. Such an enhancement of extra sex comb phenotype has also been described for molecular chaperone *Hsc4* mutant in *Drosophila* (10, 16). The PcG-like behavior of *Tcp-1η* mutants was supported by the antagonistic effect of *Tcp-1η* in *Tcp-1η/trx*^*1*^ double mutants where it suppresses the *trx*^*1*^ mutant (32) phenotype. This genetic data is substantiated by an increased expression of homeotic and non-homeotic targets of PcG, which is explained by dissociation of PC from chromatin when Tcp-1η was depleted. Additionally, de-repression of Hox genes in *Tcp-1η* RNAi flies mimics *Pc* RNAi phenotype in eye-antennal imaginal discs (41) further reinforcing the PC specific effect of *Tcp-1η* in transcriptional cellular memory.

Based on the genetic and molecular evidence presented here, it is plausible to assume that Tcp-1η is required for proper folding or stability of PC protein in cell. Since PC remains stable after Tcp-1η depletion (Fig. 4*J*) when compared with Tubulin, an obligate folding client of TRiC (25), it suggests that PC may not require Tcp-1η for its stability. This observation led us to hypothesize that Tcp-1η may facilitate the recruitment of PC on chromatin. This hypothesis explains the strong enhancement of extra sex comb phenotype and reactivation of PcG targets since mutations in *Tcp-1η* chaperonin might have reduced the chromatin-associated PC protein to maintain gene repression. Molecular chaperones usually transiently interact with their client proteins to help them fold properly and achieve a specific stable conformation thus enabling them to perform specific cellular functions (21, 47). However, Co-IP of Tcp-1η with PC as well as their co-localization at chromatin highlights a more stable interaction of Tcp-1η chaperonin with PC, similar to the previously described stable interactions between Polyhomeotic and molecular chaperones Hsc4 and Droj2 in *Drosophila* cells (16). The mechanistic link between PC and Tcp-1η may be similar to the role played by TRiC in priming HDAC3 for its nuclear localization and potential interaction with SMRT to form active SMRT-HDAC3 repression complex (27). Moreover, TRiC interacts with HDAC1/HDAC2 (29), SWI/SNF chromatin remodeling complexes (29) and acts as a checkpoint in assembly of basal transcription factor TFIID (30), highlighting the importance of TRiC at different nodes in regulating gene expression states. Molecular interaction of TRiC complex with HDACs, SWI/SNF complexes and its role in holo TFIID assembly indicates that TRiC chaperonins may have a dual role in silencing and activation of gene expression. Such a dual role of molecular chaperones in transcriptional activation (11, 12, 44) and repression (48–50) has also been reported for Hsp90.

Since TRiC chaperonins have been reported to regulate folding of translational machinery (51), we postulated that the reduced binding of PC on the chromatin in Tcp-1η knock-down cells may be due to defects in translation of PC. Contrary to this hypothesis, the total amount of PC in Tcp-1η depleted cells (Fig. 4*J*) was significantly increased as compared to control. Despite an increased amount of PC in Tcp-1η depleted cells, there was decreased association of PC at chromatin. This may be due to a large amount of PC retained in the cytosol in the absence of Tcp-1η. It would be interesting to determine composition of PC interacting proteins in wild type cells, and in cells deficient for Tcp-1η. If Tcp-1η is required for assembly of PC containing protein complexes, different PC interacting proteins will be purified in the absence of Tcp-1η. Since it is possible to assemble functional PRC1 core complex *in vitro* (52) in the absence of Tcp-1η, it suggests that Tcp-1η may have a specific effect on association of PC at chromatin. The depletion of PC, but not RNA pol-II from chromatin, in the absence of Tcp-1η further suggests that Tcp-1η has a specific role in PC mediated repression of key developmental genes. The increased expression of PcG targets in Tcp-1η depleted cells as a result of decreased association of PC and subsequent release of RNA pol-II from the paused state is in accordance with the previously reported function of PC in holding RNA pol-II in its paused state (17, 18, 53) on silent genes. While we provide evidence that PC requires Tcp-1η to maintain repression, an indirect effect of Tcp-1η on association of PC at chromatin through one of its client proteins cannot be ruled out.

The genetic and molecular interplay of TRiC at a multitude of cellular signaling pathways together with evolutionarily conserved PcG suggests that it may serve as a major hub similar to Hsp90 to maintain cellular homeostasis. The molecular interaction between Tcp-1η and PC illustrates how gene expression states can rapidly be modulated which eventually may lead to accumulation of epigenetic variation and possible phenotypic variation. Further molecular and biochemical characterization of Tcp-1η-PC nexus is required to reveal the mechanistic details about how this intricate relationship contributes to the complex interplay between genotype and phenotype.

## Materials and Methods

### Fly strains

The following fly strains were obtained from Bloomington *Drosophila* Stock Center: *Tcp-1η*^*KG01477*^, *Tcp-1η*^*KG09501*^, *Pc*^*1*^ (*Pc*^*1*^*/TM3Ser*), *Pc*^*XL5*^ (*Pc*^*XL5*^*/TM3Ser, Sb*), *trx*^*1*^ (*trx*^*1*^*/TM1*), *CCT5*^*06444*^, *CCT5*^*K06005*^. The fly strains obtained from Vienna *Drosophila* Resource Center include *Tcp-1η*^*KK101540*^, *Tcp-1η*^*GD13899*^. Moreover, fly strains used for polytene staining of transgenic flies include *P{w*^*+mC*^*UASp-Tcp-1η-MYC}, pTub-GAL4/Tb*. Additionally, fly strains used for immunostaining of polytene chromosomes after RNAi include *Tcp-1η*^*GD13899*^, and *P{Sgs3-GAL4*.*PD}*.

The fly strains used to perform eye-specific knock-down were *Tcp-1η*^*KK101540*^, *ey-GAL4*. For immunostaining of homozygous mutant embryos, *Tcp-1η*^*KG01477*^ *and Tcp-1η*^*KG09501*^ were balanced with a GFP balancer chromosome.

### Antibodies

The antibodies and their dilutions used in this study are as following: mouse anti-Abd-B (DSHB, 1A2E9) (IF 1:40), rabbit anti-PC (IF 1:20, WB 1:4000) (gift from S. Hirose), mouse anti-Myc (Santa Cruz, 9E10) (IF 1:50, WB 1:1000), mouse anti-FLAG M2 (Sigma Aldrich, F1804) (ChIP 5μl, WB 1:1000), mouse anti-Tubulin (Abcam) (WB 1:2000), mouse anti-RNA pol-II (Abcam) (ChIP 2μl, IF 1:100), mouse anti-Lamin (DSHB, ADL67.10-s) (WB 1:1000).

### Genetic Analysis

Two different alleles each of *Tcp-1η* and *CCT5* were independently crossed with two different *Pc* alleles (*Pc*^*1*^ and *Pc*^*XL5*^) at 25°C while *w*^*1118*^ flies crossed to *Pc* alleles were used as control. Males in the progeny of these crosses were scored for extra sex comb phenotype as described previously (11). Similarly, both mutants of *Tcp-1η* and *w*^*1118*^ were crossed to *trx* mutant allele (*trx*^*1*^) and males in the progeny of these crosses were analyzed and scored for abdominal pigmentation phenotype as described previously (31, 32).

### Immunohistochemistry

Transgenic flies carrying *P{w*^*+mC*^*UASp-Tcp-1η-Myc}* were generated using FemtoJet Microinjector (Eppendorf) following standard protocol (54). These transgenic flies were crossed with *pTub-GAL4/Tb* driver line to induce the expression of *Tcp-1η-Myc*. Salivary glands from *non-Tb* third instar larvae expressing *Tcp-1η-Myc* were isolated and polytene chromosomes were stained with anti-Myc and anti-PC antibodies using standard protocol (55). For embryonic staining, stage 15 embryos were dechorionated and GFP negative embryos (homozygous for the mutations) were separated under an epifluorescent stereo microscope (Nikon, C-DSS230) followed by immunostaining with Abd-B antibodies following standard protocol (55). All images were acquired using the Nikon C2 Confocal Microscope.

### Chromatin Immunoprecipitation (ChIP)

ChIP was performed from stable cell lines (see *SI Appendix*) expressing FLAG-Tcp-1η or empty vector (EV) induced with 500μM CuSO_4_ for 72hrs. Purified ChIP DNA from each reaction was quantified using quantitative real-time PCR (Applied Biosystems Inc).

Chromatin enrichment was calculated as fold change over input using ΔΔCt method (56, 57). For ChIP after knock-down of *Tcp-1η*, cells were treated with 10µg/mL of dsRNA for 5 days. Knock-down was confirmed using real-time PCR (Applied Biosystems Inc). ChIP was then performed with 1×10^7^ cells as described in *SI Appendix*. ChIP DNA was quantified using real-time PCR (Applied Biosystems Inc) and chromatin enrichment, for ChIP after knock-down of *Tcp-1η*, was calculated as percentage input using ΔΔCt method as described previously (57, 58).

### Co-Immunoprecipitation (Co-IP)

Co-IP was performed from stable cell lines (see *SI Appendix*) expressing FLAG-Tcp-1η or empty vector (EV) induced with 500μM CuSO_4_ for 72hrs. Total protein extract was incubated with anti-PC antibodies for immunoprecipitation followed by analysis on Western blot. The protocol for Co-IP is detailed in *SI Appendix*.

### Analysis of nuclear and cytosolic fractions

Cells were harvested and centrifuged at 10,000rpm for 5min at 4°C followed by lysis in Buffer A (pH7.9, 10mM HEPES, 1mM EDTA, 1mM EGTA, 100mM KCl, 1mM DTT, 0.5% NP-40 supplemented with protease and phosphatase inhibitors) for 15min on ice. After mixing gently on a vortex for 30seconds, it was centrifuged at 10,000rpm for 15min. The supernatant was collected in a fresh tube as cytosolic fraction and the pellet was resuspended in 2x SDS loading buffer and labeled as nuclear fraction. Nuclear and cytosolic fractions isolated from both Tcp-1η depleted and control cells were analyzed on Western blot with appropriate antibodies.

## Acknowledgements

We would like to thank Susumu Hirose at National Institute of Genetics (NIG) Japan, for sharing anti-PC antibody. We also thank Muhammad Sabieh Anwar for providing us access and guidance with imaging of eye phenotypes using Scanning Electron Microscope facility at Syed Babar Ali School of Science and Engineering, LUMS.

## Author Contributions

NS, JA, ZU and MT designed research; NS, JA, MHFK, MS performed research; NS, JA, AF and MT analyzed data; NS, JA, ZU and MT wrote the paper.

## References

1. Cavalli G, Paro R (1999) Epigenetic inheritance of active chromatin after removal of the main transactivator. Science (80-) 286(5441):955–958.

2. Beuchle D, Struhl G, Müller J (2001) Polycomb group proteins and heritable silencing of Drosophila Hox genes. Development 128(6):993–1004.

3. Kassis JA, Kennison JA, Tamkun JW (2017) Polycomb and trithorax group genes in drosophila. Genetics. doi:10.1534/genetics.115.185116.

4. Brand M, Nakka K, Zhu J, Dilworth FJ (2019) Polycomb/Trithorax Antagonism: Cellular Memory in Stem Cell Fate and Function. Cell Stem Cell 24(4):518–533.

5. Cao R, et al. (2002) Role of histone H3 lysine 27 methylation in polycomb-group silencing. Science (80-) 298(5595). doi:10.1126/science.1076997.

6. Czermin B, et al. (2002) Drosophila enhancer of Zeste/ESC complexes have a histone H3 methyltransferase activity that marks chromosomal Polycomb sites. Cell 111(2):185–196.

7. Müller J, et al. (2002) Histone methyltransferase activity of a Drosophila Polycomb group repressor complex. Cell 111(2):197–208.

8. Wang H, et al. (2004) Role of histone H2A ubiquitination in Polycomb silencing. Nature 431(7010). doi:10.1038/nature02985.

9. Klymenko T, Jürg M (2004) The histone methyltransferases Trithorax and Ash1 prevent transcriptional silencing by Polycomb group proteins. EMBO Rep 5(4):373– 377.

10. Mollaaghababa R, et al. (2001) Mutations in Drosophila heat shock cognate 4 are enhancers of Polycomb. Proc Natl Acad Sci U S A. doi:10.1073/pnas.061497798.

11. Tariq M, Nussbaumer U, Chen Y, Beisel C, Paro R (2009) Trithorax requires Hsp90 for maintenance of active chromatin at sites of gene expression. Proc Natl Acad Sci U S A. doi:10.1073/pnas.0809669106.

12. Sawarkar R, Sievers C, Paro R (2012) Hsp90 globally targets paused RNA polymerase to regulate gene expression in response to environmental stimuli. Cell. doi:10.1016/j.cell.2012.02.061.

13. Anderson M, et al. (2002) A new family of cyclophilins with an RNA recognition motif that interact with members of the trx/MLL protein family in Drosophila and human cells. Dev Genes Evol 212(3):107–113.

14. Nelson CJ, Santos-Rosa H, Kouzarides T (2006) Proline Isomerization of Histone H3 Regulates Lysine Methylation and Gene Expression. Cell 126(5):905–916.

15. Chen Y (2011) The immunophilin FKBP39 regulates polycomb group mediated epigenetic control in Drosophila melanogaster. Dissertation (ETH Zurich). doi:10.3929/ethz-a-006875876.

16. Wang YJ, Brock HW (2003) Polyhomeotic stably associates with molecular chaperones Hsc4 and Droj2 in Drosophila Kc1 cells. Dev Biol. doi:10.1016/S0012-1606(03)00396-8.

17. Breiling A, Turner BM, Bianchi ME, Orlando V (2001) General transcription factors bind promoters repressed by Polycomb group proteins. Nature. doi:10.1038/35088090.

18. Dellino GI, et al. (2004) Polycomb silencing blocks transcription initiation. Mol Cell. doi:10.1016/S1097-2765(04)00128-5.

19. Enderle D, et al. (2011) Polycomb preferentially targets stalled promoters of coding and noncoding transcripts. Genome Res. doi:10.1101/gr.114348.110.

20. Kuroda MI, Kang H, De S, Kassis JA (2020) Dynamic Competition of Polycomb and Trithorax in Transcriptional Programming. Annu Rev Biochem 89:235–253.

21. Gestaut D, Limatola A, Joachimiak L, Frydman J (2019) The ATP-powered gymnastics of TRiC/CCT: an asymmetric protein folding machine with a symmetric origin story. Curr Opin Struct Biol 55:50–58.

22. Won K-A, Schumacher RJ, Farr GW, Horwich AL, Reed SI (1998) Maturation of Human Cyclin E Requires the Function of Eukaryotic Chaperonin CCT. Mol Cell Biol. doi:10.1128/mcb.18.12.7584.

23. Camasses A, Bogdanova A, Shevchenko A, Zachariae W (2003) The CCT chaperonin promotes activation of the anaphase-promoting complex through the generation of functional Cdc20. Mol Cell. doi:10.1016/S1097-2765(03)00244-2.

24. Kaisari S, Sitry-Shevah D, Miniowitz-Shemtov S, Teichner A, Hershko A (2017) Role of CCT chaperonin in the disassembly of mitotic checkpoint complexes. Proc Natl Acad Sci U S A. doi:10.1073/pnas.1620451114.

25. Sternlicht H, et al. (1993) The t-complex polypeptide 1 complex is a chaperonin for tubulin and actin in vivo. Proc Natl Acad Sci U S A. doi:10.1073/pnas.90.20.9422.

26. Kim AR, Choi KW (2019) TRiC/CCT chaperonins are essential for organ growth by interacting with insulin/TOR signaling in Drosophila. Oncogene. doi:10.1038/s41388-019-0754-1.

27. Guenther MG, Yu J, Kao GD, Yen TJ, Lazar MA (2002) Assembly of the SMRT-histone deacetylase 3 repression complex requires the TCP-1 ring complex. Genes Dev. doi:10.1101/gad.1037502.

28. Wells CA, Dingus J, Hildebrandt JD (2006) Role of the chaperonin CCT/TRiC complex in G protein βγ-dimer assembly. J Biol Chem. doi:10.1074/jbc.M602409200.

29. Banks CAS, et al. (2018) Differential HDAC1/2 network analysis reveals a role for prefoldin/CCT in HDAC1/2 complex assembly. Sci Rep. doi:10.1038/s41598-018-32009-w.

30. Antonova S V., et al. (2018) Chaperonin CCT checkpoint function in basal transcription factor TFIID assembly. Nat Struct Mol Biol. doi:10.1038/s41594-018-0156-z.

31. Umer Z, et al. (2019) Genome-wide RNAi screen in Drosophila reveals Enok as a novel trithorax group regulator. Epigenetics and Chromatin. doi:10.1186/s13072-019-0301-x.

32. Ingham P, Whittle R (1980) Trithorax: A new homoeotic mutation of Drosophila melanogaster causing transformations of abdominal and thoracic imaginal segments - Putative role during embryogenesis. MGG Mol Gen Genet. doi:10.1007/BF00271751.

33. Dunn AY, Melville MW, Frydman J (2001) Review: Cellular substrates of the eukaryotic chaperonin TRiC/CCT. J Struct Biol 135(2):176–184.

34. Celniker SE, Keelan DJ, Lewis EB (1989) The molecular genetics of the bithorax complex of Drosophila: characterization of the products of the Abdominal-B domain. Genes Dev. doi:10.1101/gad.3.9.1424.

35. Delorenzi M, Bienz M (1990) Expression of Abdominal-B homeoproteins in Drosophila embryos. Development.

36. Simon J, Chiang A, Bender W (1992) Ten different Polycomb group genes are required for spatial control of the abdA and AbdB homeotic products. Development.

37. Schuettengruber B, et al. (2009) Functional anatomy of polycomb and trithorax chromatin landscapes in Drosophila embryos. PLoS Biol. doi:10.1371/journal.pbio.1000013.

38. Schwartz YB, et al. (2010) Alternative epigenetic chromatin states of polycomb target genes. PLoS Genet. doi:10.1371/journal.pgen.1000805.

39. Gibson G, Gehring WJ (1988) Head and thoracic transformations caused by ectopic expression of Antennapedia during Drosophila development. Development.

40. Plaza S, et al. (2001) Molecular basis for the inhibition of Drosophila eye development by Antennapedia. EMBO J. doi:10.1093/emboj/20.4.802.

41. Zhu J, Ordway AJ, Weber L, Buddika K, Kumar JP (2018) Polycomb group (PcG) proteins and Pax6 cooperate to inhibit in vivo reprogramming of the developing Drosophila eye. Dev. doi:10.1242/dev.160754.

42. Sawarkar R, Paro R (2013): An emerging hub of a networker. Trends Cell Biol 23(4):193–201.

43. Condelli V, et al. (2019) HSP90 Molecular Chaperones, Metabolic Rewiring, and Epigenetics: Impact on Tumor Progression and Perspective for Anticancer Therapy. Cells 8(6):532.

44. Sollars V, et al. (2003) Evidence for an epigenetic mechanism by which Hsp90 acts as a capacitor for morphological evolution. Nat Genet. doi:10.1038/ng1067.

45. Riley PD, Carroll SB, Scott MP (1987) The expression and regulation of Sex combs reduced protein in Drosophila embryos. Genes Dev. doi:10.1101/gad.1.7.716.

46. Pattatucci AM, Kaufman TC (1991) The homeotic gene Sex combs reduced of Drosophila melanogaster is differentially regulated in the embryonic and imaginal stages of development. Genetics.

47. Morán Luengo T, Mayer MP, Rüdiger SGD (2019) The Hsp70–Hsp90 Chaperone Cascade in Protein Folding. Trends Cell Biol 29(2):164–177.

48. Laskar S, Bhattacharyya MK, Shankar R, Bhattacharyya S (2011) HSP90 controls SIR2 mediated gene silencing. PLoS One. doi:10.1371/journal.pone.0023406.

49. Okazaki K, et al. (2018) RNAi-dependent heterochromatin assembly in fission yeast Schizosaccharomyces pombe requires heat-shock molecular chaperones Hsp90 and Mas5. Epigenetics and Chromatin. doi:10.1186/s13072-018-0199-8.

50. Sun L, et al. (2020) The molecular chaperone Hsp90 regulates heterochromatin assembly through stabilizing multiple complexes in fission yeast. J Cell Sci. doi:10.1242/jcs.244863.

51. Roobol A, et al. (2014) The chaperonin CCT interacts with and mediates the correct folding and activity of three subunits of translation initiation factor eIF3: B, i and h. Biochem J 458(2):213–224.

52. Francis NJ, Saurin AJ, Shao Z, Kingston RE (2001) Reconstitution of a functional core polycomb repressive complex. Mol Cell. doi:10.1016/S1097-2765(01)00316-1.

53. Lis J (1998) Promoter-associated pausing in promoter architecture and postinitiation transcriptional regulation. Cold Spring Harbor Symposia on Quantitative Biology doi:10.1101/sqb.1998.63.347.

54. Bienz M, et al. (1988) Differential regulation of Ultrabithorax in two germ layers of drosophila. Cell 53(4):567–576.

55. Sullivan WA, Ashburner M, Hawley RS (2000) Drosophila protocols. doi:10.1007/978-1-61779-034-8_15.

56. Tariq M, et al. (2003) Erasure of CpG methylation in Arabidopsis alters patterns of histone H3 methylation in heterochromatin. Proc Natl Acad Sci U S A 100(15):8823– 8827.

57. Green MR, Sambrook J (2018) Analysis and normalization of real-time polymerase chain reaction (PCR) experimental data. Cold Spring Harb Protoc 2018(10):769–777.

58. Lin X, Tirichine L, Bowler C (2012) Protocol: Chromatin immunoprecipitation (ChIP) methodology to investigate histone modifications in two model diatom species. Plant Methods 8(1):1–9.

59. Papp B, Müller J (2006) Histone trimethylation and the maintenance of transcriptional ON and OFF states by trxG and PcG proteins. Genes Dev. doi:10.1101/gad.388706.

