## Supplementary Information for "Polycomb requires Tcp-1η chaperonin for maintaining gene silencing in *Drosophila*"

### SI Appendix

#### Materials and Methods

##### dsRNA Synthesis

Specific regions used for *Tcp-1η* knock-down in cells (Fig. 2*J*) were selected from *Drosophila* RNAi Screening Center (DRSC) and their primers are detailed in *SI Appendix*, Table S1. Templates for the preparation of dsRNA were amplified by PCR from cDNA using T7-tailed oligonucleotides as primers. These templates were then used for *in vitro* transcription to synthesize dsRNA by using T7 Megascript kit following manufacturer's instructions (Ambion).

##### *Drosophila* Cell Culture

*Drosophila* S2 cells were cultured in Schneider's *Drosophila* medium (Gibco, ThermoFisher Scientific), supplemented with 10% fetal bovine serum (Gibco, ThermoFisher Scientific) and 1% penicillin-streptomycin (Gibco, ThermoFisher Scientific) at 25°C. *Drosophila* S2 cells adjusted to serum-free growth medium (D.Mel-2, Gibco, ThermoFisher Scientific) were cultured in Express Five SFM (Gibco, ThermoFisher Scientific) supplemented with 20mM GlutaMAX (Gibco, ThermoFisher Scientific) and 1% penicillin-streptomycin.

##### Western Blotting

Cells were lysed in ice-cold lysis buffer (150mM NaCl, 0.05M Tris, 1% Triton X-100) supplemented with protease inhibitors. After centrifugation at 10,000rpm, supernatant was collected in fresh tubes, mixed with 2X reducing sample buffer and boiled at 95°C for 5 minutes. The proteins were resolved on 10% SDS PAGE and transferred onto nitrocellulose membranes that were blocked with 5% milk for 30min before probing with the appropriate antibodies overnight at 4°C. Secondary antibodies, HRP conjugated, were used at 1:10,000 dilution and blots were developed using ECL reagents (GE Healthcare). The experiments were reproduced thrice.

**Generation of Stable Cell Line and Transgenic Flies**

To generate vectors expressing tagged proteins, RNA extracted from *Drosophila* S2 cells was used to synthesize cDNA. Primers designed for Gateway Cloning were used to amplify *Tcp-1η* CDS from cDNA which was then cloned into *pENTR™/D-TOPO®* vector (ThermoFisher Scientific). LR Clonase reactions (ThermoFisher Scientific) were set up with DGRC (*Drosophila* Genomics Resource Center) vectors for both cell culture and fly transformation containing either Myc or FLAG tags to prepare epitope tagged Tcp-1η. Primer details can be found in *SI Appendix,* Table S1.

For the generation of stable cell lines, *pMT-FLAG-Tcp-1η* plasmid was transfected into *Drosophila* S2 cells using Effectene transfection reagent (Qiagen). Transfected cells were selected with Hygromycin B (Roche) to a final concentration 250µg/mL. Finally, cells were induced with 500µM $CuSO_4$ for 72hrs and stable cells were confirmed for the expression of FLAG-Tcp-1η by Western blotting with anti-FLAG antibody.

Fly transformation vector expressing Myc-Tcp-1η, under the control of *UASp,* was used to generate transgenic fly lines by injecting $w^{1118}$ embryos using standard protocol (1). Transgenic flies were confirmed by Western blotting with anti-Myc antibody.

**Chromatin Immunoprecipitation (ChIP)**

ChIP was performed as described previously (2) with slight modifications. Briefly, $3x10^7$ cells were fixed at room temperature with 1% formaldehyde for 10mins. Cross-linking was stopped by the addition of glycine to a final concentration of 0.125M. Cells were washed with 1x PBS and lysed with Buffer A (10mM Tris pH 8.0, 0.25% Triton X-100, 10mM EDTA, 0.5mM EGTA) followed by two washes with Buffer B (10mM Tris pH 8.0, 200mM NaCl, 1mM EDTA, 0.5mM EGTA). Cells were sonicated in 300µl of sonication buffer (10mM Tris pH8.0, 1mM EDTA, 0.5mM EGTA) using Bioruptor (Diagenode) at high setting for 15-25mins (30sec ON, 30sec OFF) such that chromatin fragment sizes were between 100-500bp. Sonicated chromatin was centrifuged at 13,000rpm for 10mins at 4°C and the cleared chromatin was stored at -80˚C. Chromatin was diluted with 2x RIPA buffer (20mM Tris pH8.0, 2mM EDTA, 280mM NaCl, 2%Triton X-100, 0.2% SDS, 0.2% Sodium deoxycholate) and precleared by incubating with DYNA beads (Invitrogen) for 2hrs at 4˚C with 20rpm rotation. Precleared chromatin was incubated with the appropriate antibody overnight at 4˚C with 20rpm rotation. Immunocomplexes were pulled down with DYNA beads. The beads were washed 5 times with 1x RIPA, once with LiCl Buffer (10mM Tris pH8.0, 250mM LiCl, 1mM EDTA, 0.5% NP-40, 0.5% Sodium deoxycholate) and twice with 1x TE (10mM Tris pH8.0, 1mM EDTA). Chromatin was eluted by incubating beads at 65˚C with 500µl of freshly made elution buffer (0.1M Sodium bicarbonate, 1% SDS) for 15mins. Reverse crosslinking of chromatin was carried out overnight with NaCl at 65˚C followed by Proteinase K treatment for 2hrs. Reverse cross linked chromatin was extracted using phenol-chloroform followed by ethanol precipitation. All buffers were supplemented with PMSF, Aprotinin, Leupeptin and Pepstatin protease inhibitors (ThermoFisher Scientific). Primers used in ChIP analysis can be found in *SI Appendix*, Table S1.

**Co-Immunoprecipitation (Co-IP)**

Briefly, *FLAG-Tcp-1η* expressing stable cells and *EV* cells induced for 72hrs were lysed in lysis buffer (50mM NaCl, 50mM Tris pH8.0, 1mM EDTA, 0.2mM $Na_3VO_4$, 1% Triton X-100) supplemented with protease inhibitors [Pepstatin (0.5µg/ml), Leupeptin (0.5µg/ml), Aprotinin (0.5µg/ml) and PMSF (1mM)]. Lysis was carried out on ice for 10min followed by sonication (three cycles each of 5sec ON and 10sec OFF) to shear the DNA. The lysate was centrifuged at 14000rpm for 10min and supernatant was transferred to a fresh tube. The samples were quantified by Bradford reagent (ThermoFisher Scientific) and equal proteins were taken as IP samples from both *FLAG-Tcp-1η* and *EV* cells. 1% of the total lysates were used as input samples. The lysates were then incubated with anti-PC antibody at 4°C with 20rpm rotation on an orbital shaker. After 2hrs of incubation, the immune complexes were incubated with DYNA beads (Invitrogen) for 4hrs at 4°C with 20rpm rotation. The beads were then washed thrice with lysis buffer for 5min each to remove the unbound proteins. Finally, the immunoprecipitated (IP) samples together with their respective Input samples were resuspended in 2x SDS loading dye, boiled at 95°C for 5min and analyzed on a Western blot with respective antibodies.

#### Figure Legends

**Figure S1. *Tcp-1η* and *CCT5* mutants enhance extra sex comb phenotype of *Pc*.**

(A-D) $Tcp\text{-}1\eta^{KG09501}$ and $CCT5^{K06005}$ mutants were crossed to two different *Pc* ($Pc^{1}$ and $Pc^{XL5}$) alleles and double mutant (*Tcp-1η/Pc and CCT5;Pc*) male flies in the progeny were scored for extra sex comb phenotype. Heterozygous male flies for *Pc* (*+/Pc*) from the cross of $w^{1118}$ with *Pc* alleles were used as control. $Tcp\text{-}1\eta^{KG09501}$ mutation enhanced the extra sex comb phenotype of both $Pc^{1}$ and $Pc^{XL5}$ in double mutant $Tcp\text{-}1\eta^{KG09501}/Pc^{1}$ (A) and $Tcp\text{-}1\eta^{KG09501}/Pc^{XL5}$ (B) as compared to control. Similarly, $CCT5^{K06005}$ mutant showed increase in the extra sex comb phenotype in double mutant $CCT5^{K06005};Pc^{1}$ (C) and $CCT5^{K06005};Pc^{XL5}$ (D) progeny as compared to control. Severity of phenotype and statistical analysis was performed as described in Figure 1.

**Figure S2. Western blot confirms the expression of epitope-tagged *Tcp-1η* transgene.**

(A) Western blot showed Myc-tagged-Tcp-1η specifically detected in larval extracts where the expression of *UAS-Myc-Tcp-1η* was induced by crossing with pTub-GAL4 (+) driver line as compared to control (-). (B) Western blot showed the expression of FLAG-Tcp-1η in stable cells induced with 500μM $CuSO_4$ (+) as compared to un-induced control (-). Tubulin was used as loading control. (C) Headless pupal lethal pharate adult obtained as a result of eye specific knock-down of *Tcp-1η*.

**Supplementary Table**

Table S1. Primers list used in this study.

**References:**

1. Bienz M, et al. (1988) Differential regulation of Ultrabithorax in two germ layers of drosophila. *Cell* 53(4):567–576.

2. Tariq M, Nussbaumer U, Chen Y, Beisel C, Paro R (2009) Trithorax requires Hsp90 for maintenance of active chromatin at sites of gene expression. *Proc Natl Acad Sci U S A*. doi:10.1073/pnas.0809669106.

**Table S1: List of primers used in this study**

| Sr No. | Primer Name | Sequence | Purpose |
|---|---|---|---|
| 1 | T7-LacZ F | **TAATACGACTCACTATAGGGAGA**GGAAGATCAGGATATGTGG | LacZ dsRNA |
| 2 | T7-LacZ R | **TAATACGACTCACTATAGGGAGA**CTTCATCAGCAGGATATCC | |
| 3 | T7-Tcp-1η F | **TAATACGACTCACTATAGGGAGA**GCAGCCACTGCCATGTC | Tcp-1η dsRNA |
| 4 | T7-Tcp-1η R | **TAATACGACTCACTATAGGGAGA**CCACCGCAAGCCTTCAT | |
| 5 | Tcp-1η F | CACCATGCAACCGCAAATCGTGCT | Primers used for pENTR™/D-TOPO® cloning |
| 6 | Tcp-1η R (NS) | CATGGGCCTGCCCATTCCG | |
| 7 | Tcp-1η R (WS) | TTACATGGGCCTGCCCATTCCG | |
| 8 | bxd-s-low | GCACTTAAAACGGCCATTACGAA | Primers used for analysis of ChIP |
| 9 | bxd-s-up | GACGTGCGTAAGAGCGAGATACAG | |
| 10 | Dfd F | AACTCTCCGTGCGAGCGAAC | |
| 11 | Dfd R | ATGCTCCCTCTCAGTCGCGCT | |
| 12 | Intergenic Region F (IR) | CCGAACATGAGACATGGAAAA | |
| 13 | Intergenic Region R (IR) | AAAGTGCCGACAATGCAGTTA | |
| 14 | psq_TSS_F | ATAAGGCGATGCCACCTAGTTA | |
| 15 | psq_TSS_R | AATGTAGCAAAAGGTGCTCAAAG | |
| 16 | Dfd_GB1_F | ACTACTTGCAAAAGCAGCGC | |
| 17 | Dfd_GB1_R | GAAACTTTGGGTCCAAGCCAT | |
| 18 | Dfd_GB2_F | ATGGGCTCAGTTGAGTTGAC | |
| 19 | Dfd_GB2_R | TATGGTCGAACTGGAGTATC | |
| 20 | Act57B F | TGTGTGACGATGAAGTTGCTGC | Primers used for qPCR Analysis |
| 21 | Act57B R | ATCACCGACGTACGAGTCCTT | |
| 22 | Tcp-1η_RT-F | TGATTGTGGATGCCCACGG | |
| 23 | Tcp-1η_RT-R | TGGGTGCACTCCCTCCTCC | |
| 24 | AbdB ex1.1 F | CAACTACCGAACTAAGCTGC | |
| 25 | AbdB ex1.2 R | CACAATGAGGAGCAAGGATG | |
| 26 | Dfd F | CGATGGCGAACGGATCATCTA | |
| 27 | Dfd R | GCGTCAGGTAGCGGTTGTAGTGG | |
| 28 | Ubx F | ATGAACTCGTACTTTGAACAGGC | |
| 29 | Ubx R | CCAGCGAGAGAGGGAATCC | |
| 30 | Antp F | GCCTCCGCTGGTGGATCAAAT | |
| 31 | Antp R | GCTGGTACATGCCCATGTTGTGAT | |
| 32 | psq_E3F | GCAAACATCCCACAATTATCCT | |
| 33 | psq_E4R | TCGCAGAGTCCCTTGATCTT | |
| 34 | Pc_F | GGAGTAAGGGGAAGTTGGGGCG | |
| 35 | Pc_R | CGGCGATCCAGGATGTTTAC | |

#### Figure S1.

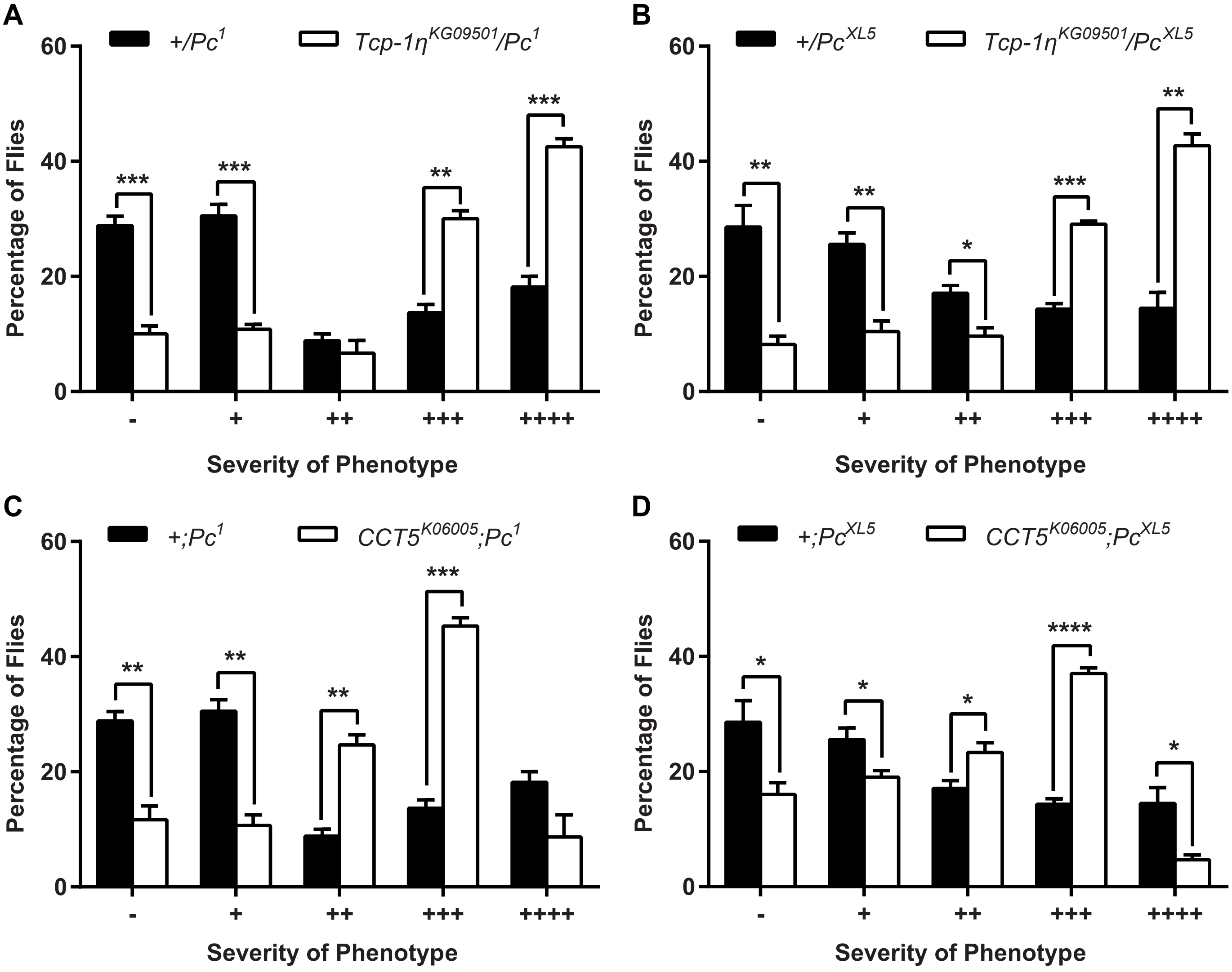
A
+/Pc1
Tcp-1ηKG09501/Pc1
Percentage of Flies
Severity of Phenotype
- + ++ +++ ++++
B
+/PcXL5
Tcp-1ηKG09501/PcXL5
Percentage of Flies
Severity of Phenotype
C
+;Pc1
CCT5K06005;Pc1
Percentage of Flies
Severity of Phenotype
D
+;PcXL5
CCT5K06005;PcXL5
Percentage of Flies
Severity of Phenotype

### Figure S2.

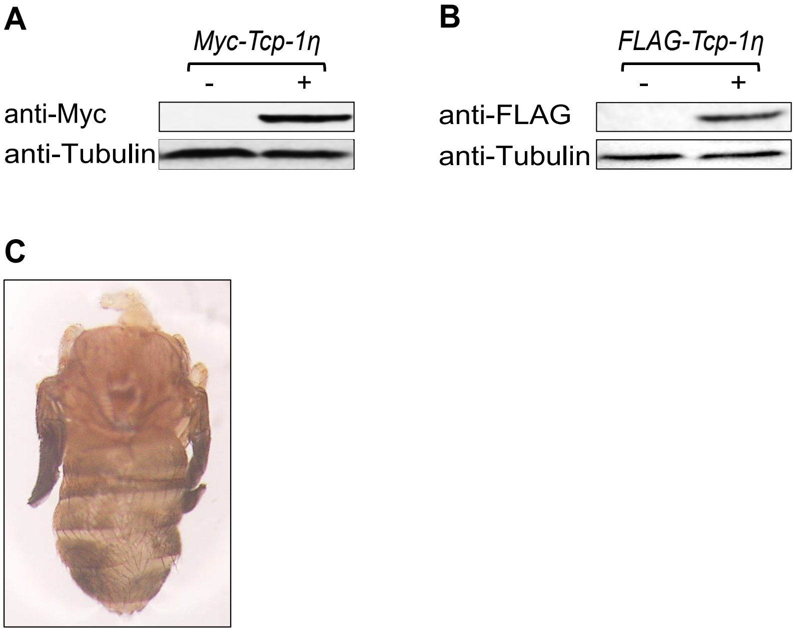
A
Myc-Tcp-1η
- +
anti-Myc
anti-Tubulin
B
FLAG-Tcp-1η
- +
anti-FLAG
anti-Tubulin
C
